# How environment and genetic architecture of unreduced gametes shape the establishment of autopolyploids

**DOI:** 10.1101/2024.09.13.612821

**Authors:** Yu Cheng, Filip Kolář, Roswitha Schmickl, Josselin Clo

## Abstract

It is broadly assumed that polyploidy success is due to an increase in fitness associated with whole genome duplication due to higher tolerance to stressful conditions. In agreement, several theoretical models found that, among other factors, a better tolerance to new environmental conditions can promote polyploidy establishment. These models, however, often made strong hypotheses, for example considering that diploids cannot adapt to new conditions, or that unreduced gametes production is not a limiting factor and that it is of a fixed quantity. In this paper, we challenged some of these hypotheses. We developed a theoretical model in which we modeled the joint evolution of a quantitative trait under selection and the production of unreduced gametes, this trait also being a quantitative trait; both traits were pleiotropically linked. We followed the adaptation of initially diploid populations to a new environment to which neo-tetraploid individuals were directly adapted. The generation of these autotetraploid individuals was enabled by the genetic production of unreduced gametes and by the environmental change modifying the average production of these gametes. We found that for realistic values of unreduced gametes production, adaptation to new environmental conditions was mainly achieved through adaptation of diploids to the new optimum rather than the fixation of newly adapted tetraploid individuals. In broader parameter sets, we found that the adaptation process led to mixed-ploidy populations, except when the populations were swamped with unreduced gametes, and that pleiotropy and environmental effects favored the co-existence of both cytotypes.

## Introduction

Polyploids are organisms known for having more than two chromosome sets compared with their diploid congener. Polyploidy resulting from whole genome duplication (WGD) is frequently accompanied by meiotic abnormalities and alters gene dosage, consequently affecting the phenotypes and long-term evolution of populations (Otto 2007). There is evidence that polyploidization is an important mechanism in speciation and diversification (Soltis *et al*. 2015), with blooms of diversity associated with ancient WGD events, both in plants, particularly the angiosperms, animals and fungi (Jiao *et al*. 2011; Van De Peer *et al*. 2017). For example, it has been suggested that all angiosperms descended from an ancient WGD event (Jiao et al., 2011; *Amborella* Genome Project, 2013), and 25 to 35% of angiosperm species have been estimated to be recently formed polyploid (i.e. neo-polyploid) species (Wood et al., 2009; Mayrose et al., 2011; Barker et al., 2016a). The geographic distribution of neo-polyploids in the angiosperms shows a latitudinal trend, with a higher frequency of polyploids at higher latitudes (Rice *et al*. 2019). It is an ongoing debate if polyploidy fosters adaptability to stressful environments (Madlung 2013), and if in particular neo-polyploidy is associated with stressful conditions, such as cold and dry conditions, as reviewed by (Van De Peer *et al*. 2021). This environmental association becomes particularly intriguing when considering the mechanisms leading to the formation of neo-polyploids.

Although somatic doubling could potentially play a role in the formation of neo-polyploids, it is currently assumed that it usually involves unreduced gametes (UG), which are gametes with the somatic ploidy level, such as diploid gametes in a diploid organism; for the cytological mechanisms and molecular regulatory networks underlying UG formation see (Brownfield & Kohler 2011; De Storme & Geelen 2013). Pathways to polyploidy can be ‘one-step’ (fusion of two unreduced gametes; rather rare because of the rarity of unreduced gamete formation), ‘triploid bridge’ (fusion of unreduced and reduced gamete within a diploid population, with triploid offspring producing unreduced gametes that backcross with the reduced gamete parent), and ‘hybrid bridge’ (same as triploid bridge, only that it refers to interspecific crosses; rather common, as unreduced gamete formation is higher in hybrids) (Mason & Pires 2015). Unreduced gamete formation has been shown to depend on abiotic and biotic factors, such as temperature, moisture, nutrition and herbivory (Kreiner *et al*. 2017a; Mason & Pires 2015; Ramsey & Schemske 1998a).

Once polyploids are formed, they face numerous challenges to finally establish in a population, and to form a distinct polyploid lineage that may eventually speciate from its diploid progenitor. Upon emergence, polyploids tend to experience a lack of mating opportunities (minority cytotype exclusion; Levin 1975). The lack of mating opportunities in short-lived neo-polyploid organisms can be theoretically overcome by a strong fitness advantage of neo-polyploids (Husband, 2004), polyploidy-associated mating system transitions (Griswold 2021; Oswald & Nuismer 2011), higher clonal reproduction of polyploids (Van Drunen & Friedman 2022), but also certain spatial components of neo-polyploid establishment (Kauai *et al*. 2023). Unreduced gametes production per se is assumed to come with an immediate fitness disadvantage, due to meiotic mishap and low triploid fitness (Kreiner *et al*. 2017), but also as a result of the genetic architecture of UG formation. In the formation of unreduced gametes, several genes have been found to be involved (Bretagnolle & Thompson 1995; Brownfield & Kohler 2011; De Storme & Geelen 2013), for which pleiotropic effects (i.e. the fact that several traits are affected by a single locus) can be assumed. Under the assumption of a correlated production of male and female unreduced gametes, polyploid fixation is more likely in combination with mating system transitions, as selfing of an individual that produces unreduced gametes of both sexes will likely result in polyploid progeny. For flowering plants, self-fertilization is indeed assumed to be more common in polyploids compared to diploids (Barringer 2007), although this study did not discriminate between autopolyploids (i.e. multiplied chromosome set within a species) and allopolyploids (i.e. multiplied chromosome set from two species), which does not allow to distinguish between the effect of WGD and hybridization. Later, Husband et al. (2008) found that autopolyploids are predominantly outcrossing, while allopolyploids are largely selfing. However, all above mentioned theoretical considerations strongly depended on key assumptions. The first one is that newly formed polyploid individuals are at least as fit as their diploid progenitors (Husband 2004; Kauai *et al*. 2023; Oswald & Nuismer 2011; Van Drunen & Friedman 2022). However, neo-polyploid lineages are generally less fit than their diploid progenitors (at least in controlled conditions, see Porturas et al., 2019; Clo and Kolář, 2021 for synthetic neo-auto/allopolyploids). Considering such an initial fitness cost strongly affects the theoretical outcome, such as the evolution toward self-fertilization being disfavored in these conditions (Clo *et al*. 2022; Gaynor *et al*. 2023). In the longer term, however, autopolyploids may exhibit a fitness advantage over their diploid progenitors (Clo, 2022a), especially in scenarios of environmental change (Van De Peer *et al*. 2021b).

The second strong assumption is that UG production is high enough for the production of neo-polyploid lineages not to be a limiting factor in polyploid evolution (Baack 2005; Husband 2004; Li *et al*. 2004; Van Drunen & Friedman 2021). However, the production of unreduced gametes is generally much lower in natural populations of non-hybrid species (between 0.1 to 2%; Kreiner et al., 2017) compared to what is found in theoretical models (Felber 1991; Husband 2004; Oswald & Nuismer 2011; Van Drunen & Friedman 2022). Few models attempted to model the effect and evolution of UG production on the probability of polyploid evolution (Clo *et al*. 2022; Gerstner *et al*. 2024). Clo *et al*. (2022) treated the evolution of unreduced gametes as a quantitative trait and found that under realistic conditions high enough levels of UG production for autopolyploidy to invade are only reached when genetic drift is strong. Gerstner *et al*. (2024) modeled the inter-generation variation of unreduced gametes following the empirical data by Kreiner *et al*. (2017b) and found that such variation can lead to the evolution of autopolyploidy in initially diploid populations. While both models are informative, they are also built on strong assumptions. Clo et al. (2022) made the genetic architecture of UG production too simplistic, omitting, for example, the fact that mutations increasing UG production are pleiotropically linked to fitness and strongly affect fitness components such as pollen and/or ovule production (d’Erfurth *et al*. 2010; Erilova *et al*. 2009; Ravi *et al*. 2008; Wang *et al*. 2010; see also Brownfield & Kohler 2011 for a review). They also neglected the effects of the environment, while such factors, like temperature, are known to have major effects on UG production rates (De Storme *et al*. 2012; Mason *et al*. 2011; Pecrix *et al*. 2011; Schindfessel *et al*. 2023; Wang *et al*. 2024). On the opposite, Gerstner *et al*. (2024) made the hypothesis that the variation in UG production is only environmental, while it is known that this trait is determined by both environmental and genetic factors (Parrott & Smith 1986; Tavoletti *et al*. 1991a). Considering a more realistic genetic architecture of UG production could strongly impact the conclusions of the above-mentioned models, and could make us understand the relative contribution of both genetic and environmental effects in UG production and polyploidy evolution.

In this paper, we investigate the effect of the genetic architecture of UG production in the origin and establishment of autopolyploids in both stable and disturbed environments, by considering both environmental stochasticity and the pleiotropic effect of mutations affecting UG production on fitness. We build an individual-based simulation model extended from Clo *et al*. (2022), in which we test the following parameters: environmental factor and pleiotropy as effects of studying how unreduced gametes as a quantitative trait evolve and affect the origin and establishment of polyploids. In conclusion, we found that for realistic values of UG production, adaptation to new environmental conditions was mainly realized by adaptation of diploids to the new optimum rather than the fixation of newly adapted tetraploid individuals. In broader parameter sets, we found that the adaptation process led to mixed-ploidy populations, except when the populations were swamped with unreduced gametes, and that pleiotropy and environmental effects favored the co-existence of both cytotypes.

## Material and methods

### General assumptions

We considered the joint evolution of a quantitative trait *Z* and the production of unreduced gametes (UG) in a population of constant size *N*, initially made of diploid individuals reproducing through obligate random mating. The phenotypic value *z* of an individual was determined by the additive action of *L*__Z__ loci each with an infinite possible number of alleles and was given by

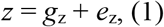

where *g*__z__ was the genetic component of the individual’s phenotype and was given by 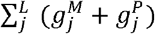, where 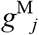 (respectively 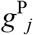) was the additive allelic effect at locus *j* inherited from the maternal (respectively paternal) gamete in the diploid population. After polyploidization and with tetrasomic inheritance, the genetic component became 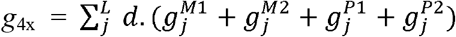, where 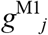 and 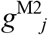 (respectively 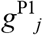 and 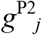) were the additive allelic effects at locus *j* inherited from the maternal (respectively paternal) gametes. The parameter *d* controlled for the dosage effect and determined the effect of polyploidization on the tetraploid genotypic values compared to diploid ones. The additive value of a new mutant allele was drawn from a Gaussian distribution of mean 0 and variance *a*^*2*^. The random environmental effect on the phenotypic trait *e*__z__ was drawn from a Gaussian distribution of mean 0 and variance *V*__E__ and was considered to be independent of the genetic components of fitness.

The trait underwent stabilizing selection around an arbitrary optimal phenotypic value, denoted *Z*__opt__ and being equal to zero in this model. The fitness value *W*__Z__ of an individual with phenotype *z* was thus described by the Gaussian function:

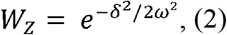

where δ was the distance between the individual’s phenotype *z* and the optimum trait value and ω^2^ was the width of the fitness function, representing the strength of selection. There was no dominance at the phenotypic scale in this model, but recessivity of mutations arose naturally at the fitness scale due to the non-linearity of the phenotype-fitness function (see Manna *et al*. 2011 for diploids and Clo, 2022 for tetraploids).

The trait “unreduced gametes production” was coded for both male and female reproductive functions by *L*__G-M__ = *L*__G-F__ freely recombining bi-allelic loci. The average UG production for male and female functions *p*__UG-M__ and *p*__UG-F__ of an individual was determined by the additive action of alleles at different loci each with an infinite possible number of alleles and was given by

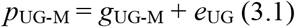

and

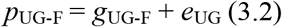

where *g*__UG-M__ (respectively *g*__UG-F__) was the genetic component of the individual’s male (respectively female) UG production and was given by 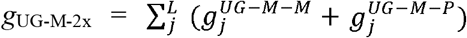, (respectively 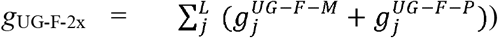, where 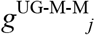 (respectively 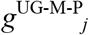) was the additive allelic effect at locus *j* inherited from the maternal (respectively paternal) gamete in the diploid population. After polyploidization and with tetrasomic inheritance, the genetic component became (for male) 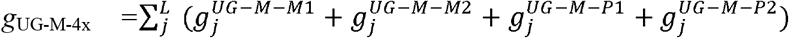, where 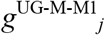 and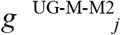 (respectively 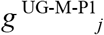 and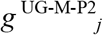) were the additive allelic effects at locus *j* inherited from the maternal (respectively paternal) gametes. The same applied for female (un)reduced gametes production. The additive value of a new mutant allele was drawn from a Gaussian distribution of mean 0 and variance *a*^*2*^. The environmental effect *e*__UG__ was fixed during simulations but could vary between the two steps of the simulation process. If by chance *p*__UG-M__ and/or *p*__UG-F__ were < 0 (respectively > 1), then *p*__UG-M__ and/or *p*__UG-F__ were forced to be equal to 0 (respectively 1). In this model, we simulated three levels of pleiotropy: 1) all the different loci of the different traits can be independent, or 2) pleiotropy between loci involved in UG production for male and female functions can happen, or 3) pleiotropy between all kind of loci (phenotypic trait and unreduced gametes for male and female functions) can occur.

### Simulation process

The simulations took place in two steps: a first one of stabilizing selection, and a second one of directional selection (adaptation process). For this step, we initiated each simulation from a population of genetically identical diploid individuals that are homozygous for the ancestral allele at all loci (*z =* 0 & *p*__UG-M__ = *p*__UG-F__ = 0 at *t* = 0). The life cycle can be summarized in five successive events. First, there is a fitness-dependent choice of the first parent (selection), followed by the choice of the second parent, which is also fitness-dependent. Selection takes place as follows: an individual is sampled randomly among the *N* individuals, but its sampling probability is weighted by its fitness. Once the two parents are chosen, the type of gametes (reduced or unreduced) they will produce is chosen. For each reproducer, one number is sampled in a uniform distribution between 0 and 1. If *p*__UG__ (male or female) is higher than the sampled value, the reproducer generates an unreduced gamete. For reduced (*n*) gametes (*n* = x and *n* = 2x gametes for diploid and tetraploid individuals, respectively), one allele per locus of each sister chromatid is sampled randomly for each trait. This phase is then followed by the introduction of mutations for each trait, the number of which is sampled from a Poisson distribution with parameter *U*, the haploid genomic mutation rate (with *U* = *µL*, where *µ* is the per-locus mutation rate and *L* is the number of loci underlying the trait under study). If the resulting offspring is not diploid or tetraploid, it will not contribute to the next generation (i.e. triploid individuals are considered inviable or sterile). The reproduction phase stops when *N* viable offspring are formed. The first step stops when the population is at Mutation-Selection-Drift (M-S-D) equilibrium. The population is considered to be at M-S-D equilibrium when the average population fitness value calculated over the last thousand generations does not differ by more than one percent from the mean fitness calculated over the previous thousand generations.

The second step of the simulations began once the population reached M-S-D equilibrium, by introducing an environmental change (done by changing the phenotypic optimum *Z*__opt__ from 0 to 2.5). After the environmental change, we assumed the polyploids are adapted to their new conditions (while before the change the fitness of each cytotype is approximately the same) and have a fitness advantage over diploids. In addition, the environmental change can induce or not a change in UG production (i.e. there is a positive non-zero effect of the environment on UG production). The population adapted either by (i) making diploid individuals to adapt to the new conditions thanks to standing genetic diversity and/or *de novo* mutations, (ii) increasing the number of newly adapted tetraploid individuals thanks to genetic diversity for unreduced gametes and the potential effect of the environment, or (iii) a mix of the two solutions (mixed-ploidy populations). The population evolved during 500 generations after the change, and the frequency and variance of unreduced gametes, the average fitness of the population, and the frequency of tetraploids in the population were inferred at each generation.

### Parameter sets

Simulations were run for several parameter sets. The values chosen for the mutation rate *U* range from 0.005 to 0.1, reflecting the per-trait haploid genomic mutation rate found in the literature (Halligan & Keightley 2009). We used parameter set values similar to those explored in similar theoretical models (Bürger *et al*. 1989; Ronce *et al*. 2009; Oswald & Nuismer 2011), with the number of freely recombining loci under selection *L* = 100, and *a*^2^ = 0.05, var(e__Z__) = var(e__UG__) = 1. The strength of stabilizing selection is set to ω^2^ = 9 to fit empirical observations (Clo & Opedal 2021). We considered a population size of *N* = 200 individuals. The dosage parameter *d* is equal to 0.65 (*g*__Z-4x__ = 1.3\**g*__Z-2x__, which is found in neo-polyploid and established polyploid populations on average; see Porturas et al., 2019 and Clo & Kolář, 2021 for meta-analyses). When pleiotropy is considered, the number of pleiotropic loci can be equal to 0 (no pleiotropy), 40 (moderate pleiotropy), or 80 (high pleiotropy). As mentioned earlier, before the environmental change, the fitness of neo-polyploids was close to the one of diploids (slightly lower because of the gigas phenotypic effect) but was initially higher at the beginning of the environmental change. Before the change, e__UG__ equals 0 but can be equal to 0, 0.1, or 0.3 after the change when considering that the environmental stress (such as change in temperature) also affects UG production. Finally, *Z*__opt__ switched from 0 to 2.5 after the environmental change. A summary of the parameters used and their values can be found in Table 1.

**Table 1.** Description of the model parameters, their abbreviations, and tested values in simulations.

| Parameter | Abbreviation | Value(s) |
| --- | --- | --- |
| Population size | $N$ | 200 |
| Number of loci | $L$ | 100 |
| Haploid genomic mutation rate | $U$ | 0.005, 0.05 or 0.1 |
| Variance of mutational effects | $a^2$ | 0.05 |
| Dosage effect | $d$ | 0.65 |
| $V_E$ of environmental effects | $V_E$ | 1 |
| Strength of stabilizing selection | $\omega^2$ | 9 |

## Results

### Description of populations at Mutation-Selection-Drift equilibrium

Under stable environmental conditions (i.e. *Z*__opt__ = 0), the main factors affecting population metrics were pleiotropy and the mutation rate (Figure 1; Supplementary Figures 1 and 2). Under a fixed mutation rate (*U* = 0.005; see below for details), pleiotropy on the quantitative trait and on UG production generally led to more efficient purging of deleterious mutations (both on the quantitative trait under stabilizing selection and on mutations increasing UG production), resulting in lower UG production (median 1% with pleiotropy vs. median 2% without pleiotropy; Figure 1A), lower genetic diversity for UG production (median 0.18% with pleiotropy vs. median 0.30% without pleiotropy; Figure 1B) and slightly increased population fitness (median 94.4% with pleiotropy vs. median 94.2% without pleiotropy; Figure 1C). Higher mutation rates (*U* = 0.05 and 0.1) did not change the patterns qualitatively, but generally led to higher average UG production (median 18% without pleiotropy for *U* = 0.05 vs. median 35% for *U* = 0.1; Supplementary Figures 1 and 2), higher genetic variance of unreduced gametes (median 18% without pleiotropy for *U* = 0.05 vs. median 36% for *U* = 0.1; Supplementary Figures 1 and 2) and lower population fitness (median 91% without pleiotropy for *U* = 0.05 vs. median 85% for *U* = 0.1; Supplementary Figures 1 and 2). Under low mutation rates, we did not find mixed-ploidy populations at the M-S-D equilibrium (0% of tetraploids, the populations were purely diploid; Figure 1D), while intermediate and high mutation rates can lead to stable mixed-ploidy populations (between 1 to 10% of tetraploids, the higher when no pleiotropic effects were modeled; Supplementary Figures 1 and 2). Because the simulations with low mutation rates gave the most realistic pattern of UG production for empirical rates of UG production (1–2%), we described separately the evolution of polyploidy with this biologically realistic scenario in the first time and other scenarios in the second time.

**Figure 1:**
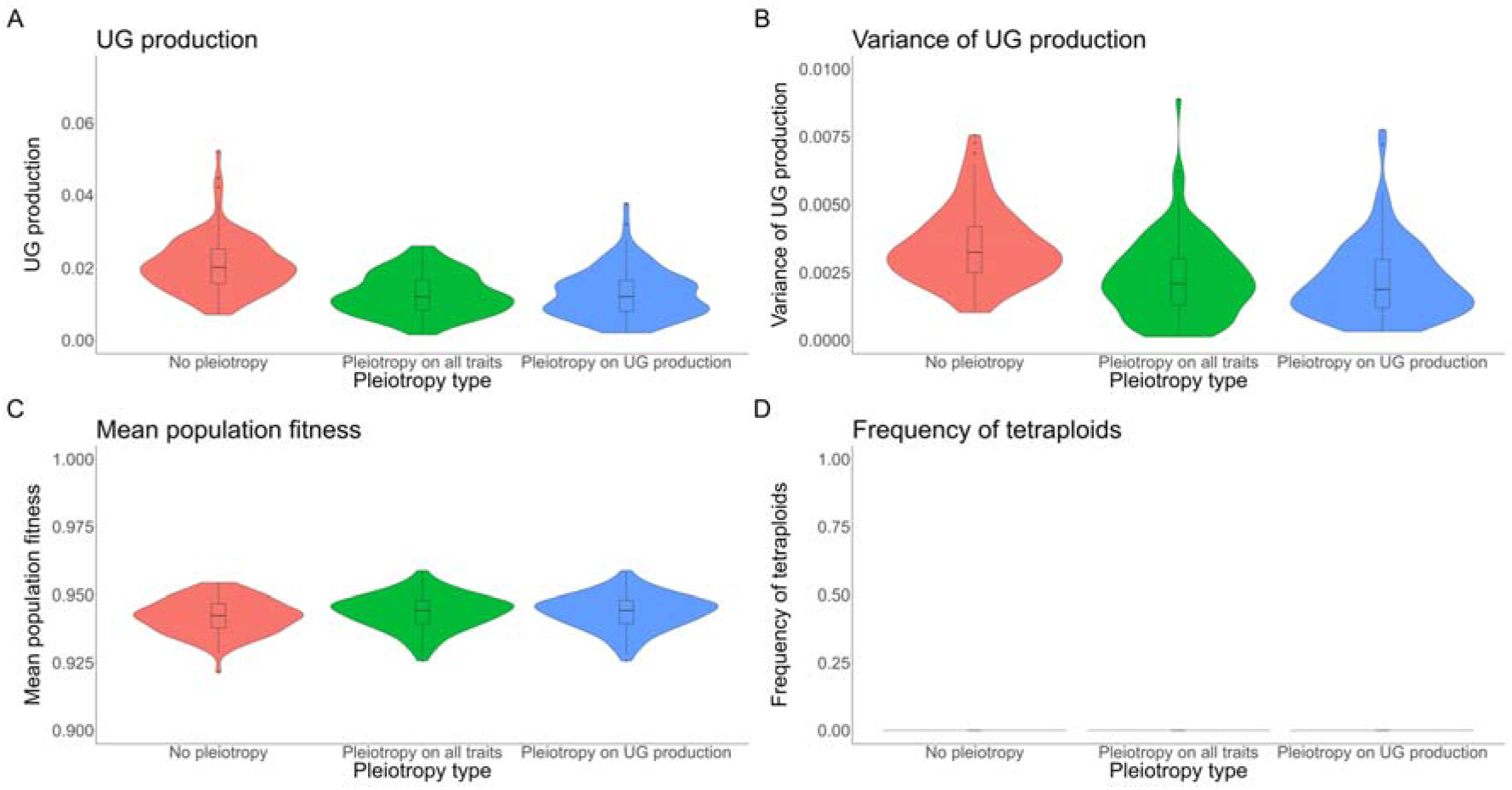
Effect of pleiotropy type (no, all traits, unreduced gametes; number of pleiotropic loci = 80) on unreduced gametes production and its variance, population fitness and tetraploid fixation, simulated from a population of genetically identical diploid individuals under the mutation rate U = 0.005. Pleiotropy on all traits means that pleiotropy affected both the quantitative trait under directional selection and unreduced gametes production. (A) Proportion of unreduced gametes production in diploid individuals; (B) variance of unreduced gametes production in diploid individuals; (C) population fitness; (D) frequency of tetraploids in the population.

### The evolution of polyploidy under environmental change for realistic parameter sets

Here we focused on simulations with low mutation rates (*U* = 0.005) and realistic environmental conditions (directional selection, *Z*__opt__ shifting from 0 to 2.5, and an environmental effect on UG production of 10%). It was observed that environmental conditions did not significantly alter the population cytotype structure and fitness after the directional selection episode (Figure 2). Both UG production and variance remained similar (UG production: median 2% with(out) environmental effect, no pleiotropy; Figure 2A; UG variance: median 0,3% with(out) environmental effect, no pleiotropy; Figure 2B). Mean population fitness also remained comparable after 500 generations when pleiotropy is absent (median 94%; Figure 2C), but with both environmental effect and pleiotropy population fitness decreased substantially, the highest being when pleiotropy only affected UG production (median 85%; Figure 2C). At the end of the simulations, the populations were only made of diploid individuals (Figure 2D), suggesting that the recovery of fitness is made by diploid individuals adapting to new conditions rather than the invasion of newly adapted tetraploids. Environmental effects on UG production rarely led to mixed-ploidy populations (Figure 2D). We attributed this to a conflict between environmental effect and new mutations that could have favored the selection of newly adapted tetraploids (but that had a negative pleiotropic effect on the trait under directional selection) to the detriment of the evolution of diploid individuals to the new optimum (Figure 3B; see Figure 3A for a comparison without environmental effect). Indeed, one can see that the production of unreduced gametes slightly increased at the beginning of the environmental change (generally doubling in the first 20 generations, shifting for example from 2% to 4% when no pleiotropy is modeled; Figure 3B). The production then decreased during the 500 generations of adaptation (Figure 3B).

**Figure 2:**
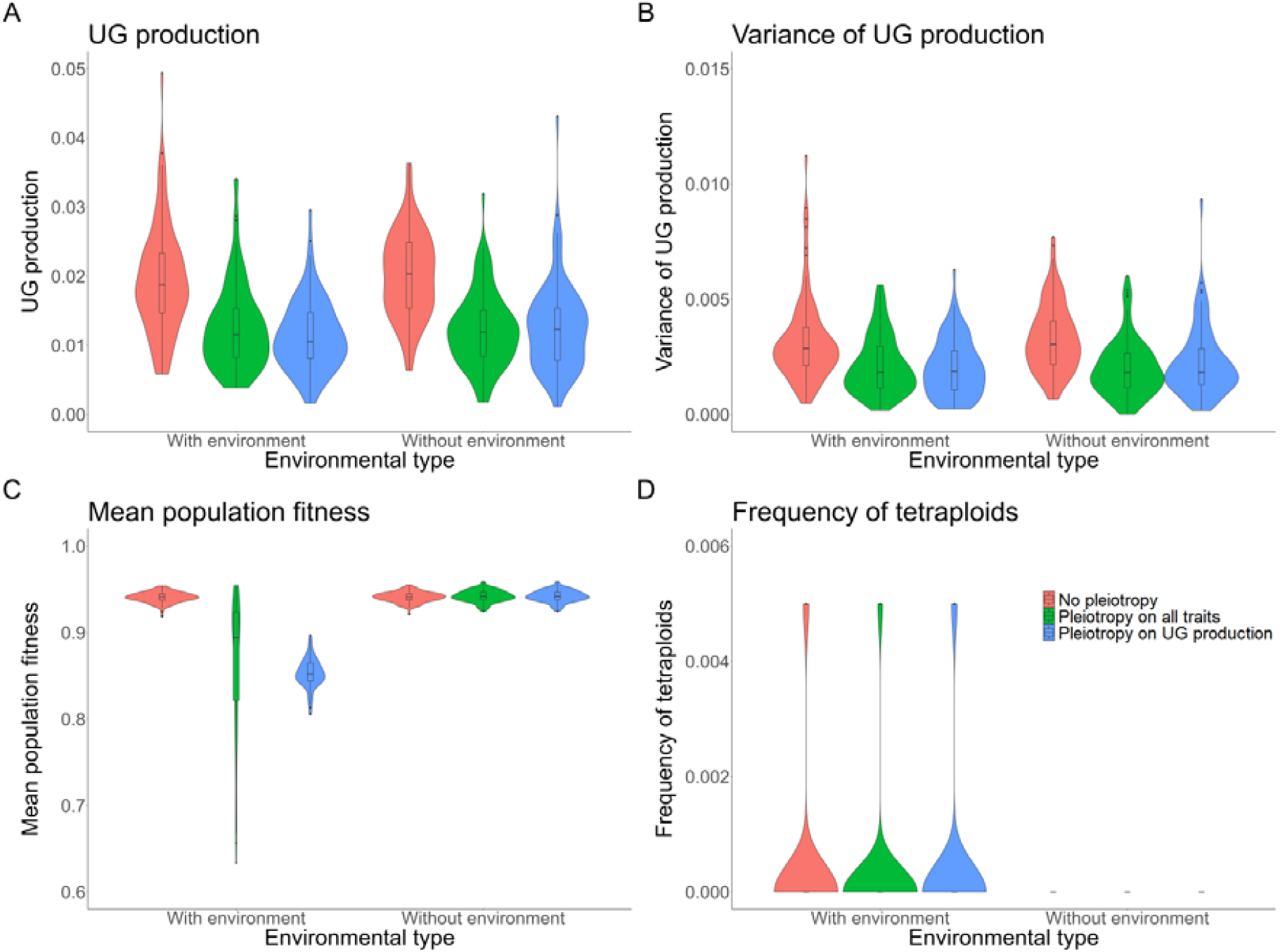
Effect of pleiotropy type (no, all traits, unreduced gametes; number of pleiotropic loci = 80) and environmental factor (0.1) on the evolution of polyploidy after 500 generations of directional selection, simulated from a population of genetically identical diploid individuals under the mutation rate U = 0.005. Pleiotropy on all traits means that pleiotropy affected both the quantitative trait under directional selection and unreduced gametes production. (A) Proportion of unreduced gametes production in diploid individuals; (B) variance of unreduced gametes production in diploid individuals; (C) population fitness; (D) frequency of tetraploids in the population.

**Figure 3:**
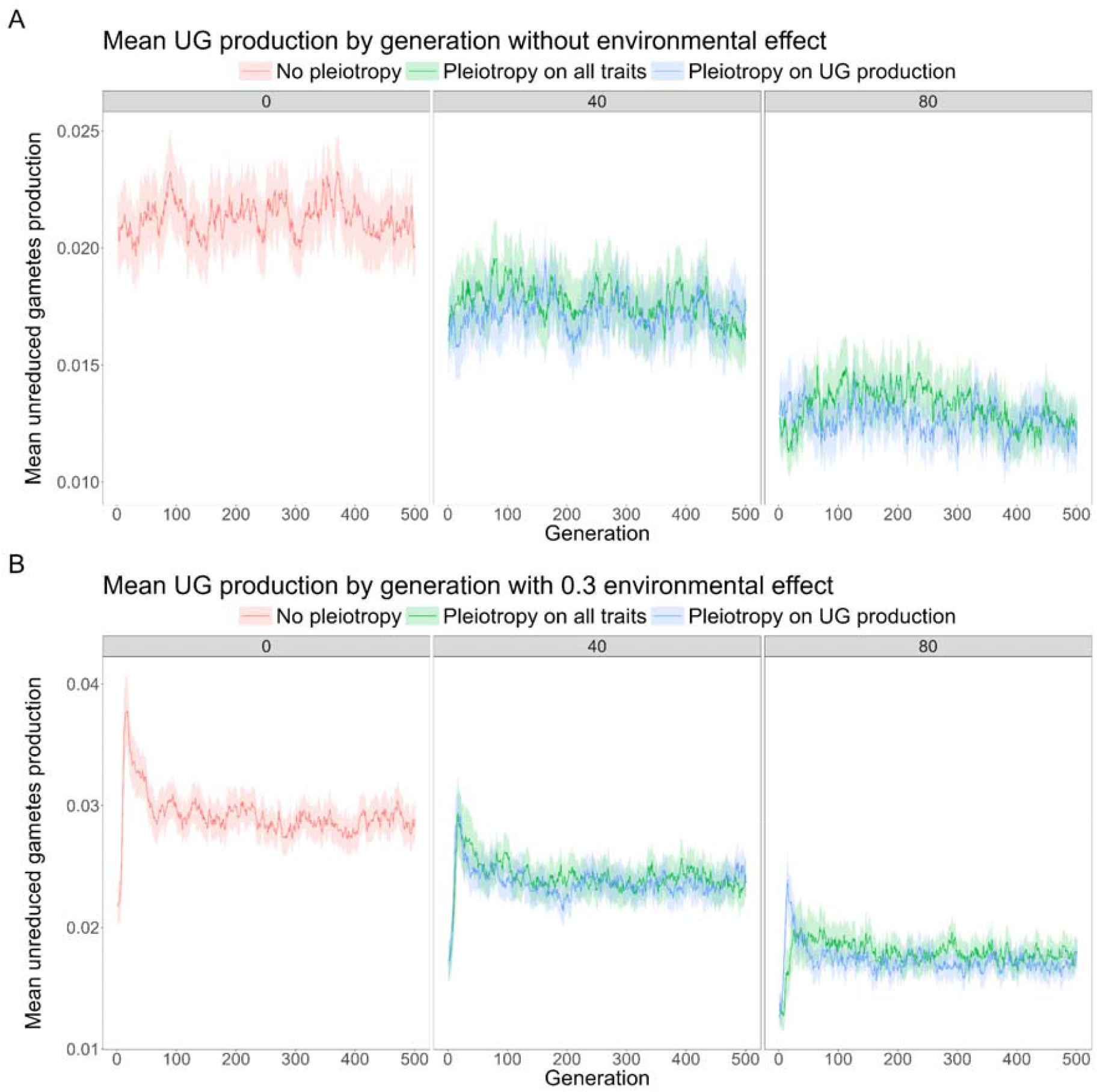
Effect of pleiotropy type (no, all traits, unreduced gametes; number of pleiotropic loci = 0, 40, 80) on unreduced gametes production (A) without effect of environmental factor; (B) with effect of environmental factor (0.3), simulated from a population of genetically identical diploid individuals under the mutation rate U = 0.005 for 500 generations. Pleiotropy on all traits means that pleiotropy affected both the quantitative trait under directional selection and unreduced gametes production.

### The general effects of environment and genetic architecture on the evolution of polyploidy

At the beginning of the directional selection episode, the frequency of unreduced gametes, due to genetic effects, increased in the first generations and then decreased back to an equilibrium value (Figures 3B and (4). This steep initial increase was due to the fact that at the beginning of the adaptation process, selection favored both the formation and the increase in frequency of newly adapted tetraploid individuals. Once diploid individuals were closer to the new phenotypic optimum, formation of tetraploids was either selected against and tetraploids remained in the minority within the population due to minority cytotype exclusion (between 0 to 10% of tetraploids when no environmental effect was modeled; Figure 5A; between 0 to 55% of tetraploids with a realistic environmental effect of 0.1 on UG production; Figure 5B), or tetraploid formation was continuously favored until fixation of tetraploidy (100% of tetraploids when the effect of the environment on UG production is strong, independent of the mutation rate; Figure 5C). As previously described, the overall effect of increased mutation rates was to increase both the average production of unreduced gametes and of tetraploids within populations (Figures 4 and 5). It was, however, notable that the mutation rate has a synergistic effect with the environmental effect on unreduced gametes: the higher the environmental effect, the higher the genetic response to increase UG production (Figure 4). This is because the production of unreduced gametes is under frequency dependence: the higher UG production is in the population, the less it is counter-selected. Thus, the stronger the effect of the environment, the easier it is for a mutation increasing UG production to be maintained in the population. Pleiotropy had a generally negative effect by decreasing the frequency of unreduced gametes and tetraploids (Figures 4 and 5). Interestingly, the effect of pleiotropy on limiting both UG production and the frequency of tetraploids was stronger when pleiotropy was limited to UG production of both male and female functions, rather than affecting both UG production and the quantitative trait under directional selection (Figures 4B and (5B). This was due to the fact that if the mutation affecting both UG production and the phenotypic trait had a positive effect on fitness (i.e. the mutation brought the phenotype closer to the new optimum), the mutation was selected despite its costly effect on UG production.

**Figure 4:**
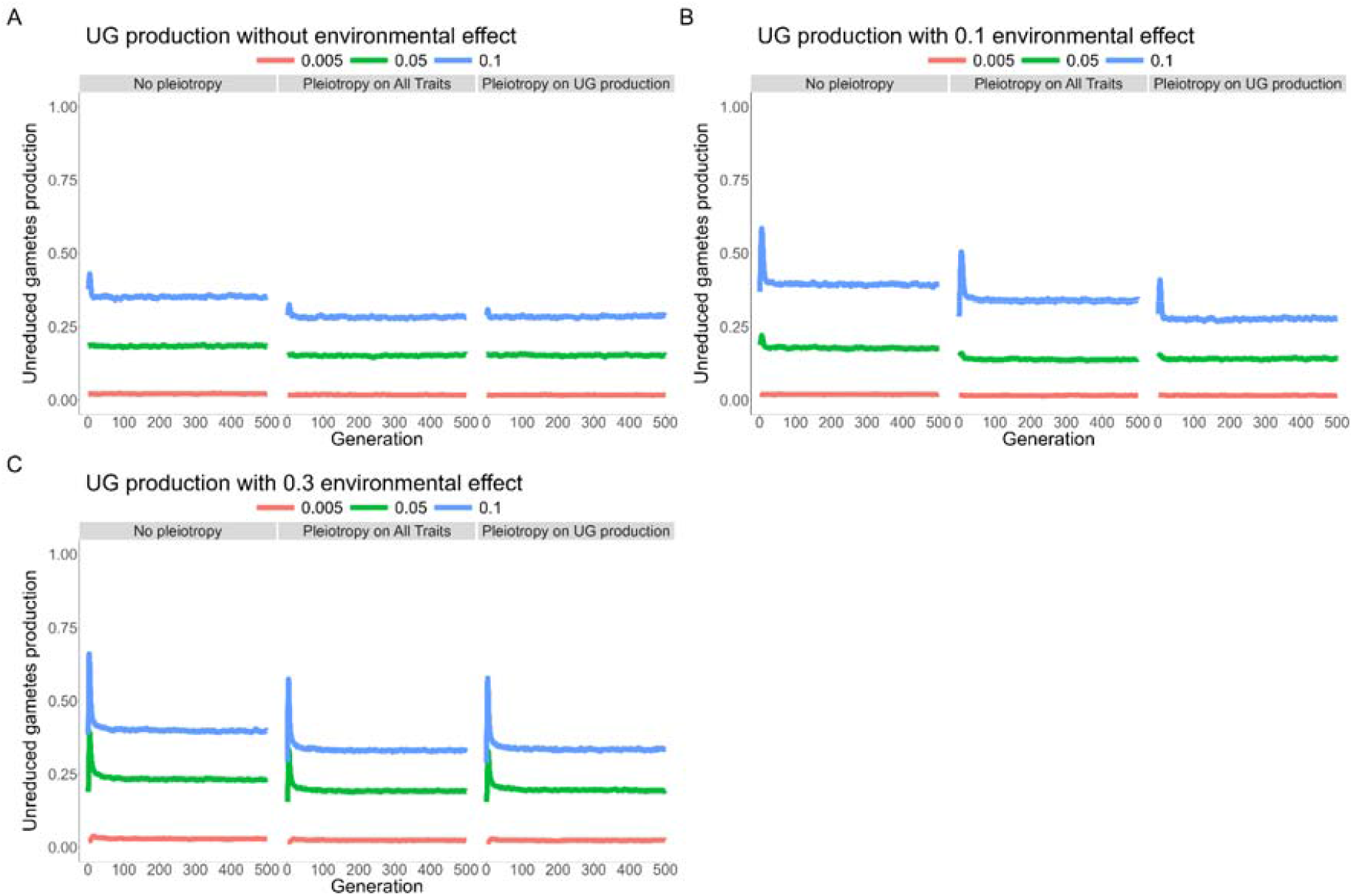
Effect of pleiotropy type (no, all traits, unreduced gametes; number of pleiotropic loci = 40) and the environmental factor (0, 0.1, 0.3) on unreduced gametes production under different mutation rates (U = 0.005, 0.05, 0.1; differently colored lines), simulated from a population of genetically identical diploid individuals for 500 generations. Pleiotropy on all traits means that pleiotropy affected both the quantitative trait under directional selection and unreduced gametes production. (A) No environmental factor; (B) environmental factor 0.1; (C) environmental factor 0.3.

**Figure 5:**
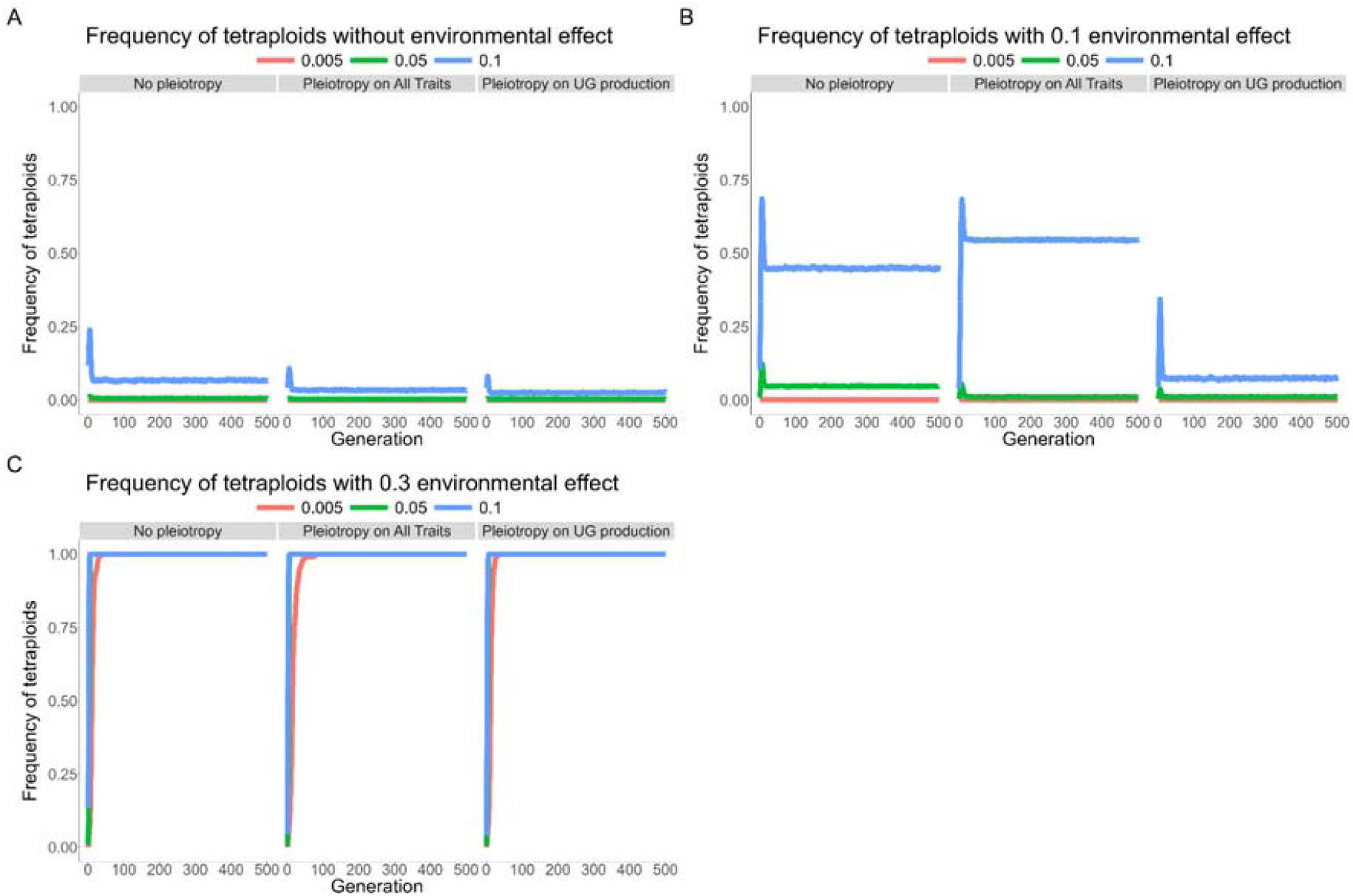
Effect of pleiotropy type (no, all traits, unreduced gametes; number of pleiotropic loci = 40) and the environmental factor (0, 0.1, 0.3) on the frequency of tetraploids in the population under different mutation rates (U = 0.005, 0.05, 0.1; differently colored lines). simulated from a population of genetically identical diploid individuals for 500 generations. Pleiotropy on all traits means that pleiotropy affected both the quantitative trait under directional selection and unreduced gametes production. (A) No environmental factor; (B) environmental factor 0.1; (C) environmental factor 0.3.

Overall, the fixation of polyploidy remained rare (Figures 5 and 6). With low mutation rates, populations adapted to new conditions either by remaining diploid, with the individuals adapting to the new conditions, or by forming mixed-ploidy populations of adapted diploids and tetraploids, notably when the effect of the environment on UG formation is high (Figures 5 and 6). With higher mutation rates, the formation of a mixed-ploidy population was much more frequent, with tetraploids reaching 10 to 55% in the populations (Figure 5B). The fixation of polyploidy was frequent when unreduced gametes swamped the population, i.e. with a high mutation rate and a strong effect of the environment (Figures 5C and (6).

**Figure 6:**
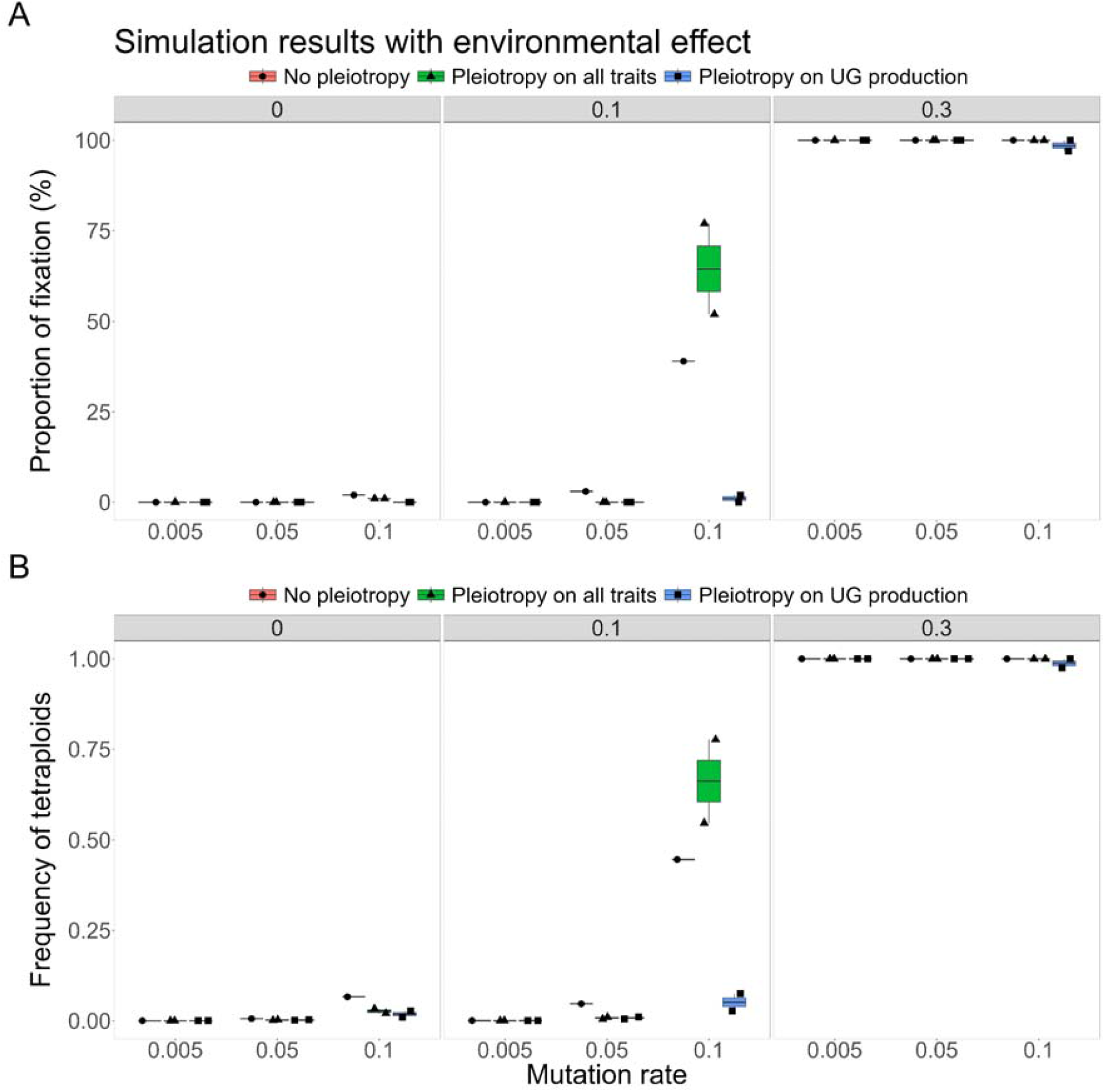
Effect of pleiotropy type (no, all traits, unreduced gametes; number of pleiotropic loci = 40) and the environmental factor (0, 0.1, 0.3) on the (A) fixation and (B) frequency of tetraploids in the population under different mutation rates (U = 0.005, 0.05, 0.1; differently colored lines). simulated from a population of genetically identical diploid individuals for 500 generations. Pleiotropy on all traits means that pleiotropy affected both the quantitative trait under directional selection and unreduced gametes production.

## Discussion & Conclusion

We simulated the interplay between stabilizing selection and directional selection in response to an environmental factor during the origin and fixation of autopolyploids. In this process, the pleiotropic effect of mutations on UG production and the quantitative trait affecting population fitness were tested. Under stabilizing selection, pleiotropy had a negative effect on the production and genetic variation of unreduced gametes. Under a realistic low mutation rate in the focal genes, we found no fixation of tetraploidy. Higher mutation rates led to higher heritable rates of UG production and to the formation of stable mixed-ploidy populations, however, with tetraploids remaining rare (< 10% in frequency). Under directional selection, the environmental factor had a complex effect on UG production and population cytotype structure. Taken as an isolated parameter, the higher the effect of the environment on UG production, the higher the frequency of polyploidy at the end of the 500 simulated generations following the environmental change. In addition, environment and genetic architecture had a synergistic effect: the stronger the effect of the environment, the higher the genetic component of UG production. This is mainly because the production of unreduced gametes is frequency-dependent, i.e. the higher it is, the less costly it is.

### Evolution of tetraploidy during stressful conditions

Environmental stress is considered to be important to the origin of polyploids, as it affects UG production (Kreiner *et al*. 2017a; Mason & Pires 2015; Ramsey & Schemske 1998a). In particular, high and low temperatures have been shown to impact male UG formation (De Storme *et al*. 2012; Mason *et al*. 2011; Pecrix *et al*. 2011; Schindfessel *et al*. 2023; Wang *et al*. 2017), although the plants used therein were mainly of hybrid origin, which makes them generally more prone to form unreduced gametes (Ramsey & Schemske 1998a). Treatment with high temperatures is commonly used to generate unreduced male gametes for polyploid breeding (Mai *et al*. 2019). In our simulations, tetraploids got fixed in a population either under extreme environmental effects or through higher genetic mutation rates that increase UG production to ca. 20 to 40% at equilibrium (*U* = 0.05, 0.1; Figure 5C; Supplementary Figure 2). These results are in line with the recent Gerstner et al. (2024) model which found that environmental stochasticity for UG production can lead to the evolution of polyploidy. However, as UG production in natural populations of non-hybrid species is typically much lower, ranging from 0.1 to 2% (Kreiner *et al*. 2017), our high mutation rates seem very unrealistic. If we assumed a realistic mutation rate for an average of 2% of UG production, tetraploids only got fixed in a population under a strong environmental effect in our simulations, while a moderate environmental effect resulted in a mixed-ploidy population with rare tetraploids (< 10%; Figure 5C). However, we found that the higher the effect of the environment on UG production, the higher the genetic increase of UG formation. If the environmental stress also affects the mutation rate (like U.V. exposure that could affect both temperature and alter the mutation rate; Gengyo-Ando & Mitani 2000), it is possible that higher rates of polyploids could be found.

In the first simulated generations after the onset of the directional selection episode, which represents the environmental change, we observed increased UG production and an increased frequency of newly adapted tetraploid individuals in the population compared to later generations (Figure 4). Under directional selection, mutations leading to UG formation might be beneficial for tetraploid establishment, while counteracting diploid adaptation. Polyploids may initially benefit from phenotypic shifts associated with WGD (Clo & Kolář 2021; Porturas *et al*. 2019). However, both of these meta-analyses are based on studies of established natural populations, which prevents filtering out the effects of selection and genetic drift that are known to have shaped population dynamics of established polyploids, but have less affected the first generations after a WGD event. The immediate, short-term effect of a WGD can only be assessed using experimentally induced, synthetic autopolyploids. For example, Westermann *et al*. (2024) showed that pollen tube tip growth is drastically challenged in immediate response to a WGD event: in synthetic autotetraploids of *Arabidopsis arenosa* and *Arabidopsis thaliana* pollen tubes grew slowly and abnormally, while in established tetraploids of *A. arenosa* pollen tube physiology and morphology returned to normal. This suggests initially reduced polyploid formation and fitness shortly after a WGD event - a scenario not modeled here. Empirical frequencies of neo-polyploids in natural populations might thus be lower. Decreased polyploid origin and fitness may also stem from the formation of triploid offspring, which have low viability due to the triploid block (Burton & Husband 2000a; Ramsey & Schemske 1998b). However, as we did not account for triploid formation in our model, this does not affect the simulations.

Our findings are in contrast to that of (Clo 2022b) in that the author found neo-tetraploidy to be associated with an initial decrease in their adaptive potential compared to their diploid progenitors, while within less than 100 generations of the simulation they reached or exceeded the additive variance of the diploids. In our simulations, tetraploids were directly adapted and selected for an initial around 50 generations, but then were selected against once diploid individuals were closer to the new phenotypic optimum, and they remained in the minority within populations, resulting in mixed-ploidy populations under moderate environmental effects. Even if the link between polyploidy and adaptive potential is probably highly multifactorial (Clo 2022a), modeling non-adapted tetraploids with smaller genetic diversity might have favored the diploid cytotype in mixed-ploidy populations.

Especially for plants, mixed-ploidy populations have been frequently reported in multiple diploid-autotetraploid species, with varying proportions of diploids and their polyploid derivatives (Kolář *et al*. 2017). The strength of selection against polyploid cytotypes in these populations further depends on the mating system and clonal propagation, although their interactions are not fully understood, based on both empirical data (Barringer 2007; Hojsgaard 2018; Van Drunen & Husband 2019) and theoretical modeling (Griswold 2021; Oswald & Nuismer 2011; Van Drunen & Friedman 2022). Eventually, gene flow between ploidy levels may transfer novel adaptive alleles (e.g., in *A. arenosa*, Baduel *et al*. 2018; Australian burrowing frogs of the genus *Neobatrachus*, Novikova *et al*. 2020) and may contribute to tetraploid fixation in the long-term, but we did not model this scenario. In our simulations, only under a strong environmental effect tetraploids came to fixation. This is in line with the theoretical framework of Oswald & Nuismer (2011) who showed that neo-tetraploid populations can adapt more efficiently than their diploid progenitors if the populations undergo an environmental change, and that this change is aligned with the phenotypic modifications found in tetraploid individuals. The duplication of entire genomes immediately creates redundant informational ‘modules’ in the genome, offering possibilities for a wider exploration of genotypic and phenotypic space (Yao *et al*. 2019). It also supports the hypothesis that polyploids are expected to face higher extinction rates than their diploid progenitors (Levin 2019) in the absence of environmental change, due to their smaller geographical ranges upon emergence. In contrast, polyploids may thrive in stressful environments and broaden their ranges, leading to their prevalence in such environments and during historical periods of climate change (Bomblies 2020; Rice *et al*. 2019; Van De Peer *et al*. 2021b). In sum, while our model did not include post-WGD environmental adaptation, it shows that environmental change may support polyploid establishment by directly affecting the rate of UG formation, moreover, with a further positive feedback effect on its underlying genes.

### The consequences of UG genetic architecture on polyploidy evolution

The production of unreduced gametes is genetically determined (Parrott & Smith 1986; Tavoletti *et al*. 1991b). It varies among individuals within and between populations, such that most individuals in natural populations of non-hybrid species produce male unreduced gametes at zero or low frequency and only a small number of individuals produce substantially more (Kreiner *et al*. 2017a). Unreduced gamete mutations affect various cytological stages (Bretagnolle & Thompson 1995; Brownfield & Kohler 2011; De Storme & Geelen 2013), and they involve pre-meiotic, meiotic and post-meiotic pathways. Key proteins comprise DYAD, AtPS1, JAS, TES, MPK4 (meiotic) and INCENP, RBR (post-meiotic). Some of these genes are polygenic (Bretagnolle & Thompson 1995; Brownfield & Kohler 2011; De Storme & Geelen 2013), which may account for the variation in UG frequencies in natural populations and the variance of UG production found in our simulations. A clear theoretical and empirical framework for the effect of genetic architecture of UG production, in particular pleiotropic effects, on the fixation of polyploidy is lacking. It is unclear if a few major effect mutations or numerous small effect mutations are involved, and how they affect the further trajectory of polyploid adaptive evolution.

In our simulations, one of the main findings is that pleiotropy always has a negative effect on UG production and its variance. For example, a higher number of pleiotropic loci significantly reduces UG production (Figure 3). This indicates that pleiotropy constrains UG production by increasing the cost of complexity, a general effect of pleiotropy (Bomblies & Peichel 2022; Orr 2000). While UG production and variance under pleiotropic effects is lowered during stabilizing selection in our simulations, the overall fitness of the population is often increased and thus seems to benefit from pleiotropic effects (Figure 1). However, we found that during directional selection, if there is pleiotropy between UG formation and fitness-related loci, a mutation having both a positive effect for diploid adaptation and increasing UG production can be temporarily selected. It is known that mutations affecting UG formation in plants are often linked to male and/or female fitness (Brownfield & Kohler 2011). If this kind of pleiotropic mutations can increase in frequency during an environmental change, as explored by us, polyploidy could evolve by chance at least at low frequency, highlighting once again the role of stochasticity in polyploid evolution (Clo *et al*. 2022; Gerstner *et al*. 2024; Kauai *et al*. 2023).

According to empirical data, UG production is a deleterious phenotype, maintained at a low frequency in the population (Kreiner *et al*. 2017b). In general, pleiotropy on quantitative traits can reduce the selection efficacy in natural populations (Fraïsse *et al*. 2019; Zhang 2023). Consequently, in our simulations individuals with UG production faced strong purifying selection but were still maintained in the population. Pleiotropy is discussed to have an antagonistic effect between fitness and reproduction, in accordance with the theory of pleiotropy constraints on adaptive traits (Hughes & Leips 2017). Antagonistic pleiotropy or the trade-off between traits has also been found for reproduction versus aging/longevity (Austad & Hoffman 2018; Wu *et al*. 2022). Empirical work has shown that intermediate pleiotropy is a driver of adaptation in *A. thaliana* (Frachon *et al*. 2017). Besides, through evolution experiments of yeast (*Saccharomyces cerevisiae*), pleiotropy resulting from a changing environment contributed to molecular adaptation (Chen & Zhang 2020). In our simulations, environment and genetic architecture have a synergistic effect: with both an environmental effect and pleiotropy, population fitness decreased. Similarly, under environmental change, higher pleiotropic effects on single cell morphology in yeast were observed compared to a stable environment (Geiler-Samerotte *et al*. 2020). Reduction of population fitness resulting from pleiotropy could eventually lead to a reduction of the population size, and the population could then be more affected by genetic drift. This scenario would be in accordance with previous models showing that the fixation of polyploids relies on strong genetic drift more than on natural selection (Clo *et al*. 2022), favoring polyploids under environmental change.

### Limitations of the model

If our model is informative, we had to avoid some genetic and ecological aspects related to ploidy for the model to remain understandable. A first simplification is that we hypothesized triploid and higher polyploid levels to be non-viable. Triploids may play a crucial role in mixed-ploidy population dynamics (Husband 2004; Kauai *et al*. 2024). However, triploids are typically very rare in natural populations of autopolyploids, possibly being abundant only during the earliest stage of polyploid formation in a population (Burton & Husband 2000b; Husband & Sabara 2004; Kolář *et al*. 2017; Ramsey & Schemske 1998a). Also, we decided to model the fact that polyploidy is adaptive in the short-term, which can be true in some conditions (Jiang *et al*. 2022; Maherali *et al*. 2009; Wang *et al*. 2019; Wu *et al*. 2019), but not in general (Clo & Kolář 2021; Porturas *et al*. 2019). Considering equivalent fitness for diploids and tetraploids, or a decrease in fitness in polyploids, would not have drastically changed the results. If the population is swamped with unreduced gametes (high mutation rate and strong environmental effect), tetraploids would fix in a population independently of their fitness (see Clo *et al*. 2022 for an example). However, for most simulations for which we found mixed-ploidy populations, it is likely that with no fitness advantage the populations would have remained diploid most of the time.

We modeled obligately outcrossing populations, while any form of assortative mating or spatial structure favoring mating within cytotypes could increase the likelihood of fixation of polyploidy (Otto & Whitton 2000). The first reason is that the aim of our model was to simulate a more realistic architecture of UG production, and that this novelty should not modify the outcomes of previous models (Husband 2004; Van Drunen & Friedman 2022). Second, association between ploidy and different forms of assortative mating remains controversial (Clo 2022b). If the association between clonality and polyploidy is widespread (Van Drunen & Husband 2019), no clear patterns emerge for the effect of ploidy on the evolution of selfing (Husband *et al*. 2008) or plant-pollinator interactions (Rezende *et al*. 2020). Finally, we did not model demographical events following the environmental change. One of our major results is that, for realistic simulations (with UG production fitting empirical data), the adaptation of populations to new conditions was more likely by diploid individuals reaching the new phenotypic optimum, rather than by adapted tetraploids reaching fixation. We could have simulated fluctuating population size, with low-fitness individuals (diploids in our case) going extinct before reaching the reproductive stage for example. With a lower population size, any increase in tetraploid frequency would have lowered the strength of frequency-dependent selection more efficiently than in our simulations, possibly leading to mixed-ploidy or tetraploid populations more often.

## Supporting information

all supplementary figures

## Data availability

Simulation code is available at https://github.com/JosselinCLO/Model_UG_pleiotropy_environment.

## Funding

This research was funded by the Czech Science Foundation (project 22-29078K to FK). JC is supported by the CNRS. Financial support also came from the long-term research development project no. RVO 67985939 of the Czech Academy of Sciences.

## Acknowledgments

Computational resources were supplied by the project “e-Infrastruktura CZ” (e-INFRA LM2018140) provided within the program Projects of Large Research, Development and Innovations Infrastructures.

## Notes

https://github.com/JosselinCLO/Model_UG_pleiotropy_environment

### Competing Interest Statement

The authors have declared no competing interest.

## Reference

Austad, S.N. & Hoffman, J.M. (2018). Is antagonistic pleiotropy ubiquitous in aging biology? Evolution, Medicine, and Public Health, 2018, 287–294.

Baack, E.J. (2005). To succeed globally, disperse locally: effects of local pollen and seed dispersal on tetraploid establishment. Heredity, 94, 538–546.

Baduel, P., Hunter, B., Yeola, S. & Bomblies, K. (2018). Genetic basis and evolution of rapid cycling in railway populations of tetraploid Arabidopsis arenosa. PLoS Genet, 14, e1007510.

Barringer, B.C. (2007). Polyploidy and selfLfertilization in flowering plants. American J of Botany, 94, 1527–1533.

Bomblies, K. (2020). When everything changes at once: finding a new normal after genome duplication. Proc. R. Soc. B., 287, 20202154.

Bomblies, K. & Peichel, C.L. (2022). Genetics of adaptation. Proc. Natl. Acad. Sci. U.S.A., 119, e2122152119.

Bretagnolle, F. & Thompson, J.D. (1995). Gametes with the somatic chromosome number: mechanisms of their formation and role in the evolution of autopolyploid plants. New Phytologist, 129, 1–22.

Brownfield, L. & Kohler, C. (2011). Unreduced gamete formation in plants: mechanisms and prospects. Journal of Experimental Botany, 62, 1659–1668.

Bürger, R., Wagner, G.P. & Stettinger, F. (1989). How much heritable variation can be maintained in finite populations by mutation–selection balance? Evolution, 43, 1748–1766.

Burton, T.L. & Husband, B. (2000a). Fitness differences among diploids, tetraploids, and their triploid progeny in Chamerion angustifolium: mechanisms of inviability and implications for polyploid evolution. Evolution, 54, 1182–1191.

Burton, T.L. & Husband, BrianC. (2000b). FITNESS DIFFERENCES AMONG DIPLOIDS, TETRAPLOIDS, AND THEIR TRIPLOID PROGENY IN CHAMERION ANGUSTIFOLIUM: MECHANISMS OF INVIABILITY AND IMPLICATIONS FOR POLYPLOID EVOLUTION. Evolution, 54, 1182–1191.

čertner, M., Sudová, R., Weiser, M., Suda, J. & Kolář, F. (2019). PloidyLaltered phenotype interacts with local environment and may enhance polyploid establishment in Knautia serpentinicola (Caprifoliaceae). New Phytologist, 221, 1117–1127.

Chen, P. & Zhang, J. (2020). Antagonistic pleiotropy conceals molecular adaptations in changing environments. Nat Ecol Evol, 4, 461–469.

Clo, J. (2022a). Polyploidization: Consequences of genome doubling on the evolutionary potential of populations. American Journal of Botany, 109, 1213–1220.

Clo, J. (2022b). The evolution of the additive variance of a trait under stabilizing selection after autopolyploidization. J of Evolutionary Biology, 35, 891–897.

Clo, J. & Kolář, F. (2021). ShortL and longLterm consequences of genome doubling: a metaLanalysis. American J of Botany, 108, 2315–2322.

Clo, J. & Opedal, Ø.H. (2021). Genetics of quantitative traits with dominance under stabilizing and directional selection in partially selfing species. Evolution, 75, 1920–1935.

Clo, J., PadillaLGarcía, N. & Kolář, F. (2022). Polyploidization as an opportunistic mutation: The role of unreduced gametes formation and genetic drift in polyploid establishment. J of Evolutionary Biology, 35, 1099–1109.

De Storme, N., Copenhaver, G.P. & Geelen, D. (2012). Production of Diploid Male Gametes in Arabidopsis by Cold-Induced Destabilization of Postmeiotic Radial Microtubule Arrays. Plant Physiology, 160, 1808–1826.

De Storme, N. & Geelen, D. (2013). Sexual polyploidization in plants – cytological mechanisms and molecular regulation. New Phytologist, 198, 670–684.

d’Erfurth, I., Cromer, L., Jolivet, S., Girard, C., Horlow, C., Sun, Y., et al. (2010). The CYCLIN-A CYCA1;2/TAM Is Required for the Meiosis I to Meiosis II Transition and Cooperates with OSD1 for the Prophase to First Meiotic Division Transition. PLoS Genet, 6, e1000989.

Erilova, A., Brownfield, L., Exner, V., Rosa, M., Twell, D., Scheid, O.M., et al. (2009). Imprinting of the Polycomb Group Gene MEDEA Serves as a Ploidy Sensor in Arabidopsis. PLoS Genet, 5, e1000663.

Felber, F. (1991). Establishment of a tetraploid cytotype in a diploid population: Effect of relative fitness of the cytotypes. J of Evolutionary Biology, 4, 195–207.

Frachon, L., Libourel, C., Villoutreix, R., Carrère, S., Glorieux, C., Huard-Chauveau, C., et al. (2017). Intermediate degrees of synergistic pleiotropy drive adaptive evolution in ecological time. Nat Ecol Evol, 1, 1551–1561.

Fraïsse, C., Puixeu Sala, G. & Vicoso, B. (2019). Pleiotropy Modulates the Efficacy of Selection in Drosophila melanogaster. Molecular Biology and Evolution, 36, 500–515.

Gaynor, M.L., Kortessis, N., Soltis, D.E., Soltis, P.S. & Ponciano, J.M. (2023). Dynamics of mixed-ploidy populations under demographic and environmental stochasticities. BioRxiv, 2023.03. 29.534764.

Geiler-Samerotte, K.A., Li, S., Lazaris, C., Taylor, A., Ziv, N., Ramjeawan, C., et al. (2020). Extent and context dependence of pleiotropy revealed by high-throughput single-cell phenotyping. PLoS Biol, 18, e3000836.

Gengyo-Ando, K. & Mitani, S. (2000). Characterization of Mutations Induced by Ethyl Methanesulfonate, UV, and Trimethylpsoralen in the Nematode Caenorhabditis elegans. Biochemical and Biophysical Research Communications, 269, 64–69.

Gerstner, B.P., Wearing, H.J. & Whitney, K.D. (2024). Why so many polyploids? Accounting for environmental stochasticity in unreduced gamete formation lowers the perceived barriers to polyploid establishment. Journal of Evolutionary Biology, 37, 325–335.

Griswold, C.K. (2021). The effects of migration load, selfing, inbreeding depression, and the genetics of adaptation on autotetraploid versus diploid establishment in peripheral habitats. Evolution, 75, 39–55.

Halligan, D.L. & Keightley, P.D. (2009). Spontaneous mutation accumulation studies in evolutionary genetics. Annual Review of Ecology, Evolution, and Systematics, 40, 151–172.

Hojsgaard, D. (2018). Transient Activation of Apomixis in Sexual Neotriploids May Retain Genomically Altered States and Enhance Polyploid Establishment. Front. Plant Sci., 9, 230.

Hughes, K.A. & Leips, J. (2017). Pleiotropy, constraint, and modularity in the evolution of life histories: insights from genomic analyses. Annals of the New York Academy of Sciences, 1389, 76–91.

Husband, B.C. (2004). The role of triploid hybrids in the evolutionary dynamics of mixedploidy populations: TRIPLOIDS IN MIXED-PLOIDY POPULATIONS. Biological Journal of the Linnean Society, 82, 537–546.

Husband, B.C., Ozimec, B., Martin, S.L. & Pollock, L. (2008). Mating Consequences of Polyploid Evolution in Flowering Plants: Current Trends and Insights from Synthetic Polyploids. International Journal of Plant Sciences, 169, 195–206.

Husband, B.C. & Sabara, H.A. (2004). Reproductive isolation between autotetraploids and their diploid progenitors in fireweed, Chamerion angustifolium (Onagraceae). New Phytologist, 161, 703–713.

Jiang, J., Yang, N., Li, L., Qin, G., Ren, K., Wang, H., et al. (2022). Tetraploidy in Citrus wilsonii Enhances Drought Tolerance via Synergistic Regulation of Photosynthesis, Phosphorylation, and Hormonal Changes. Front. Plant Sci., 13, 875011.

Jiao, Y., Wickett, N.J., Ayyampalayam, S., Chanderbali, A.S., Landherr, L., Ralph, P.E., et al. (2011). Ancestral polyploidy in seed plants and angiosperms. Nature, 473, 97–100.

Kauai, F., Bafort, Q., Mortier, F., Van Montagu, M., Bonte, D. & Van De Peer, Y. (2024). Interspecific transfer of genetic information through polyploid bridges. Proc. Natl. Acad. Sci. U.S.A., 121, e2400018121.

Kauai, F., Mortier, F., Milosavljevic, S., Van De Peer, Y. & Bonte, D. (2023). Neutral processes underlying the macro eco-evolutionary dynamics of mixed-ploidy systems. Proc. R. Soc. B., 290, 20222456.

Kolář, F., čertner, M., Suda, J., Schönswetter, P. & Husband, B.C. (2017). Mixed-Ploidy Species: Progress and Opportunities in Polyploid Research. Trends in Plant Science, 22, 1041–1055.

Kreiner, J.M., Kron, P. & Husband, B.C. (2017a). Evolutionary Dynamics of Unreduced Gametes. Trends in Genetics, 33, 583–593.

Kreiner, J.M., Kron, P. & Husband, B.C. (2017b). Frequency and maintenance of unreduced gametes in natural plant populations: associations with reproductive mode, life history and genome size. New Phytologist, 214, 879–889.

Levin, D.A. (1975). MINORITY CYTOTYPE EXCLUSION IN LOCAL PLANT POPULATIONS. TAXON, 24, 35–43.

Levin, D.A. (2019). Why polyploid exceptionalism is not accompanied by reduced extinction rates. Plant Syst Evol, 305, 1–11.

Li, B. LH., Xu, X. LM. & Ridout, M.S. (2004). Modelling the establishment and spread of autotetraploid plants in a spatially heterogeneous environment. J of Evolutionary Biology, 17, 562–573.

Madlung, A. (2013). Polyploidy and its effect on evolutionary success: old questions revisited with new tools. Heredity, 110, 99–104.

Maherali, H., Walden, A.E. & Husband, B.C. (2009). Genome duplication and the evolution of physiological responses to water stress. New Phytologist, 184, 721–731.

Mai, Y., Li, H., Suo, Y., Fu, J., Sun, P., Han, W., et al. (2019). High temperature treatment generates unreduced pollen in persimmon (Diospyros kaki Thunb.). Scientia Horticulturae, 258, 108774.

Manna, F., Martin, G. & Lenormand, T. (2011). Fitness landscapes: an alternative theory for the dominance of mutation. Genetics, 189, 923–937.

Mason, A.S., Nelson, M.N., Yan, G. & Cowling, W.A. (2011). Production of viable male unreduced gametes in Brassica interspecific hybrids is genotype specific and stimulated by cold temperatures. BMC Plant Biol, 11, 103.

Mason, A.S. & Pires, J.C. (2015). Unreduced gametes: meiotic mishap or evolutionary mechanism? Trends in Genetics, 31, 5–10.

Novikova, P.Yu., Brennan, I.G., Booker, W., Mahony, M., Doughty, P., Lemmon, A.R., et al. (2020). Polyploidy breaks speciation barriers in Australian burrowing frogs Neobatrachus. PLoS Genet, 16, e1008769.

Oberlander, K.C., Dreyer, L.L., Goldblatt, P., Suda, J. & Linder, H.P. (2016). SpeciesLrich and polyploidLpoor: Insights into the evolutionary role of wholeLgenome duplication from the Cape flora biodiversity hotspot. American Journal of Botany, 103, 1336–1347.

Orr, H.A. (2000). Adaptation and The Cost of Complexity. Evolution, 54, 13–20.

Oswald, B.P. & Nuismer, S.L. (2011). A Unified Model of Autopolyploid Establishment and Evolution. The American Naturalist, 178, 687–700.

Otto, S.P. (2007). The Evolutionary Consequences of Polyploidy. Cell, 131, 452–462.

Otto, S.P. & Whitton, J. (2000). Polyploid incidence and evolution. Annual review of genetics, 34, 401–437.

Parrott, W. & Smith, R. (1986). Recurrent selection for 2n pollen formation in red clover 1. Crop Science, 26, 1132–1135.

Pecrix, Y., Rallo, G., Folzer, H., Cigna, M., Gudin, S. & Le Bris, M. (2011). Polyploidization mechanisms: temperature environment can induce diploid gamete formation in Rosa sp. Journal of Experimental Botany, 62, 3587–3597.

Porturas, L.D., Anneberg, T.J., Curé, A.E., Wang, S., Althoff, D.M. & Segraves, K.A. (2019). A metaLanalysis of whole genome duplication and the effects on flowering traits in plants. American J of Botany, 106, 469–476.

Ramsey, J. & Schemske, D.W. (1998a). PATHWAYS, MECHANISMS, AND RATES OF POLYPLOID FORMATION IN FLOWERING PLANTS. Annu. Rev. Ecol. Syst., 29, 467–501.

Ramsey, J. & Schemske, D.W. (1998b). Pathways, mechanisms, and rates of polyploid formation in flowering plants. Annual review of ecology and systematics, 29, 467–501.

Ravi, M., Marimuthu, M.P. & Siddiqi, I. (2008). Gamete formation without meiosis in Arabidopsis. Nature, 451, 1121–1124.

Rezende, L., Suzigan, J., Amorim, F.W. & Moraes, A.P. (2020). Can plant hybridization and polyploidy lead to pollinator shift? Acta Bot. Bras., 34, 229–242.

Rice, A., Šmarda, P., Novosolov, M., Drori, M., Glick, L., Sabath, N., et al. (2019). The global biogeography of polyploid plants. Nat Ecol Evol, 3, 265–273.

Ronce, O., Shaw, F.H., Rousset, F. & Shaw, R.G. (2009). Is inbreeding depression lower in maladapted populations? A quantitative genetics model. Evolution, 63, 1807–1819.

Schindfessel, C., De Storme, N., Trinh, H.K. & Geelen, D. (2023). Asynapsis and meiotic restitution in tomato male meiosis induced by heat stress. Front. Plant Sci., 14, 1210092.

Soltis, P.S., Marchant, D.B., Van De Peer, Y. & Soltis, D.E. (2015). Polyploidy and genome evolution in plants. Current Opinion in Genetics & Development, 35, 119–125.

Tavoletti, S., Mariani, A. & Veronesi, F. (1991a). Cytological Analysis of MacroL and Microsporogenesis of a Diploid Alfalfa Clone Producing Male and Female 2 n Gametes. Crop Science, 31, 1258–1263.

Tavoletti, S., Mariani, A. & Veronesi, F. (1991b). Phenotypic recurrent selection for 2n pollen and 2n egg production in diploid alfalfa. Euphytica, 57, 97–102.

Trávníček, P., Dočkalová, Z., Rosenbaumová, R., Kubátová, B., Szeląg, Z. & Chrtek, J. (2011). Bridging global and microregional scales: ploidy distribution in Pilosella echioides (Asteraceae) in central Europe. Annals of Botany, 107, 443–454.

Van De Peer, Y., Ashman, T.-L., Soltis, P.S. & Soltis, D.E. (2021a). Polyploidy: an evolutionary and ecological force in stressful times. The Plant Cell, 33, 11–26.

Van De Peer, Y., Ashman, T.-L., Soltis, P.S. & Soltis, D.E. (2021b). Polyploidy: an evolutionary and ecological force in stressful times. The Plant Cell, 33, 11–26.

Van De Peer, Y., Mizrachi, E. & Marchal, K. (2017). The evolutionary significance of polyploidy. Nat Rev Genet, 18, 411–424.

Van Drunen, W.E. & Friedman, J. (2022). Autopolyploid establishment depends on lifeLhistory strategy and the mating outcomes of clonal architecture. Evolution, 76, 1953–1970.

Van Drunen, W.E. & Husband, B.C. (2019). Evolutionary associations between polyploidy, clonal reproduction, and perenniality in the angiosperms. New Phytologist, 224, 1266–1277.

Wang, C., Ge, W., Yin, H., Zhang, Y. & Li, J. (2024). MicroRNAs and their targets involved in unreduced pollen formation induced by heat stress in Camellia nitidissima. Scientia Horticulturae, 323, 112462.

Wang, C.-J., Zhang, L.-Q., Dai, S.-F., Zheng, Y.-L., Zhang, H.-G. & Liu, D.-C. (2010). Formation of unreduced gametes is impeded by homologous chromosome pairing in tetraploid Triticum turgidum × Aegilops tauschii hybrids. Euphytica, 175, 323–329.

Wang, J., Li, D., Shang, F. & Kang, X. (2017). High temperature-induced production of unreduced pollen and its cytological effects in Populus. Scientific Reports, 7, 5281.

Wang, Z., Zhao, Z., Fan, G., Dong, Y., Deng, M., Xu, E., et al. (2019). A comparison of the transcriptomes between diploid and autotetraploid Paulownia fortunei under salt stress. Physiol Mol Biol Plants, 25, 1–11.

Westermann, J., Srikant, T., Gonzalo, A., Tan, H.S. & Bomblies, K. (2024). Defective pollen tube tip growth induces neo-polyploid infertility. Science, 383, eadh0755.

Wu, D., Wang, Z., Huang, J., Huang, L., Zhang, S., Zhao, R., et al. (2022). An antagonistic pleiotropic gene regulates the reproduction and longevity tradeoff. Proc. Natl. Acad. Sci. U.S.A., 119, e2120311119.

Wu, G.-Q., Lin, L.-Y., Jiao, Q. & Li, S.-J. (2019). Tetraploid exhibits more tolerant to salinity than diploid in sugar beet (Beta vulgaris L.). Acta Physiol Plant, 41, 52.

Yao, Y., Carretero-Paulet, L. & Van De Peer, Y. (2019). Using digital organisms to study the evolutionary consequences of whole genome duplication and polyploidy. PLoS ONE, 14, e0220257.

Zhang, J. (2023). Patterns and Evolutionary Consequences of Pleiotropy. Annu. Rev. Ecol. Evol. Syst., 54, 1–19.

