## Supplementary material for "How environment and genetic architecture of unreduced gametes shape the establishment of autopolyploids": all supplementary figures

Supplementary data

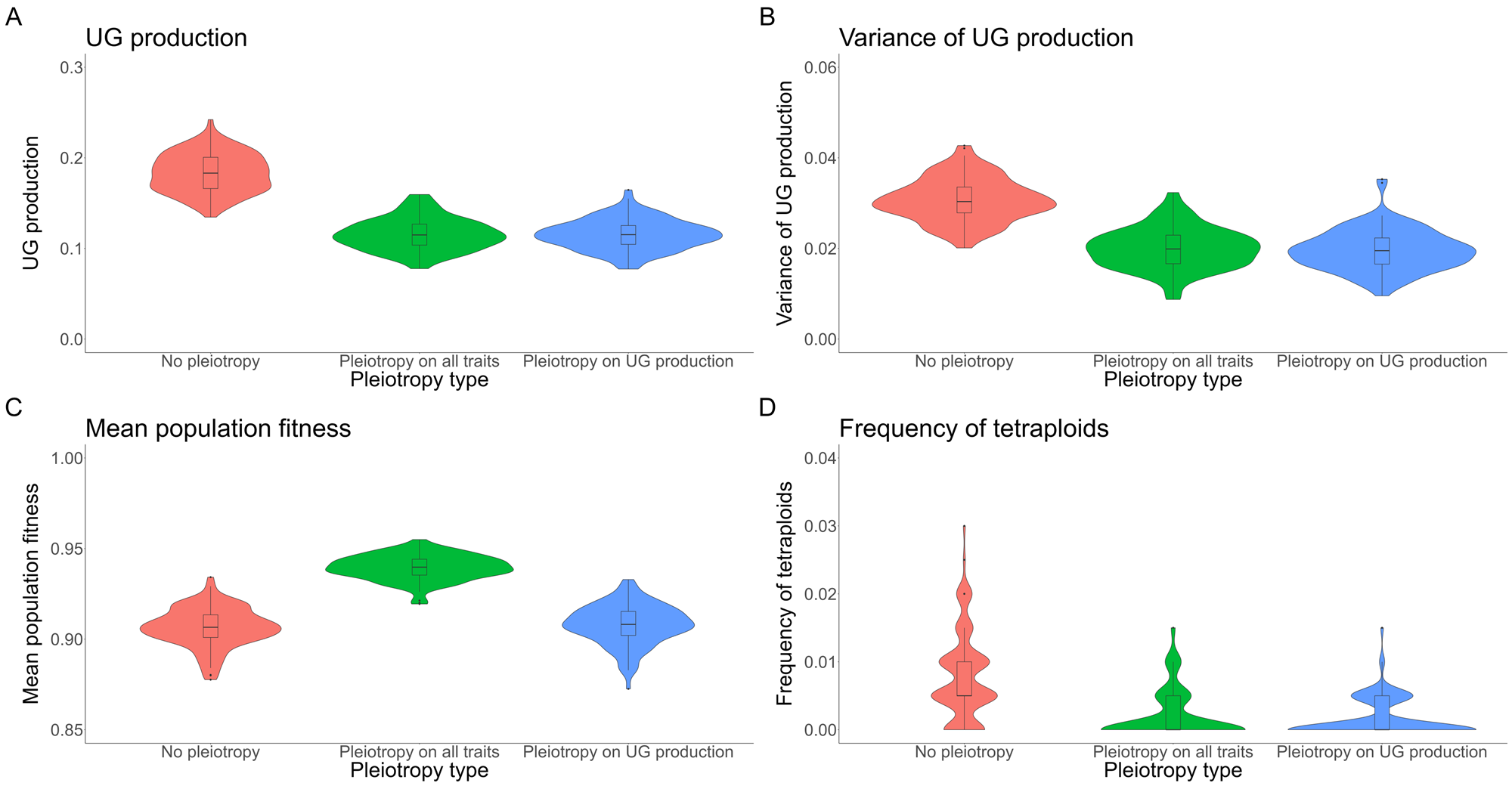

*Figure S1: Effect of pleiotropy type (no, all traits, unreduced gametes; number of pleiotropic loci = 80) on unreduced gametes production and its variance, population fitness and tetraploid fixation, simulated from a population of genetically identical diploid individuals under the mutation rate U = 0.05. Pleiotropy on all traits means that pleiotropy affected both the quantitative trait under directional selection and unreduced gametes production. (A) Proportion of unreduced gametes production in diploid individuals; (B) variance of unreduced gametes production in diploid individuals; (C) population fitness; (D) frequency of tetraploids in the population.*

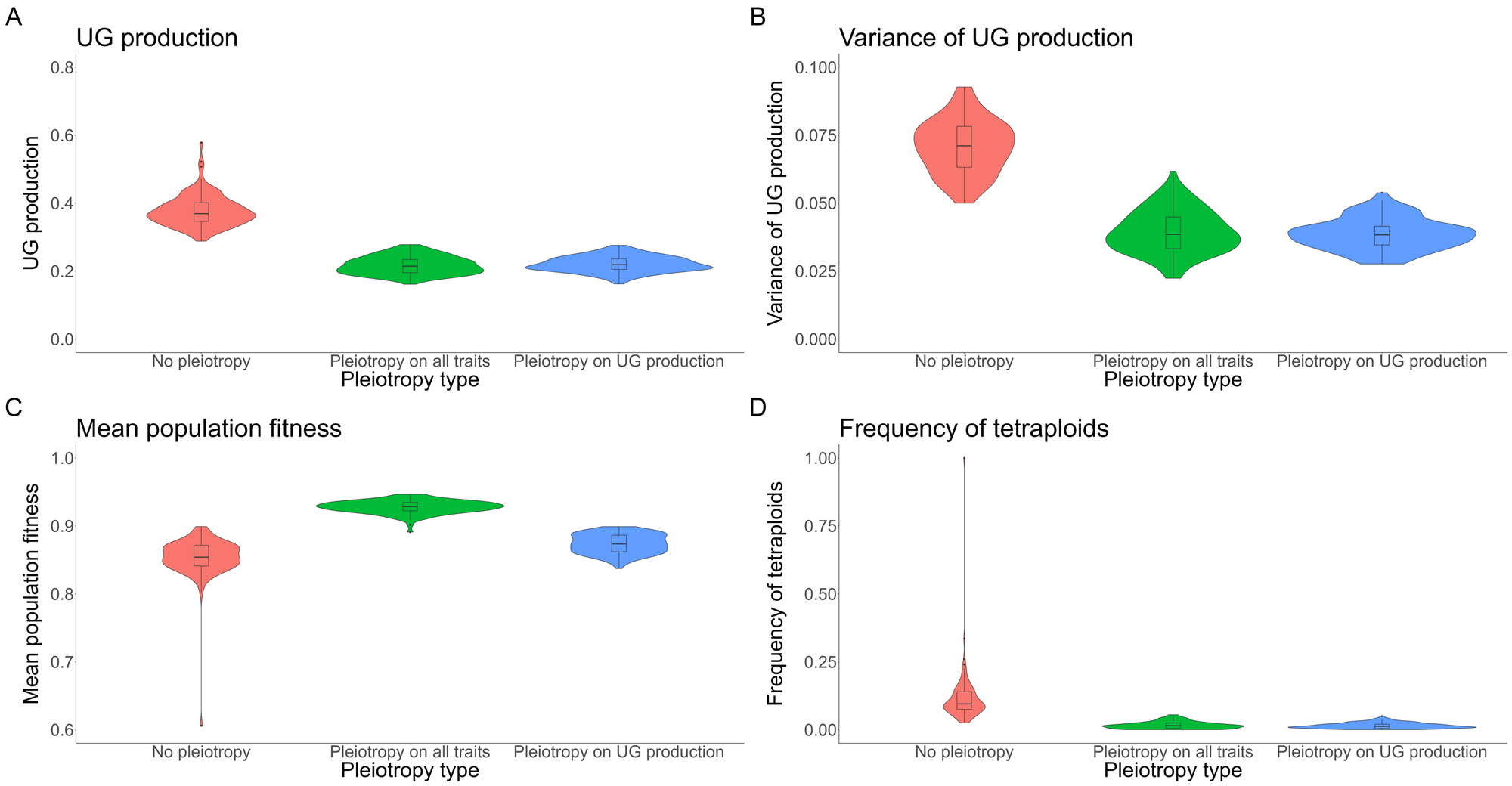

*Figure S2: Effect of pleiotropy type (no, all traits, unreduced gametes; number of pleiotropic loci = 80) on unreduced gametes production and its variance, population fitness and tetraploid fixation, simulated from a population of genetically identical diploid individuals under the mutation rate U = 0.1. Pleiotropy on all traits means that pleiotropy affected both the quantitative trait under directional selection and unreduced gametes production. (A) Proportion of unreduced gametes production in diploid individuals; (B) variance of unreduced gametes production in diploid individuals; (C) population fitness; (D) frequency of tetraploids in the population.*

**figure under environmental conditions**

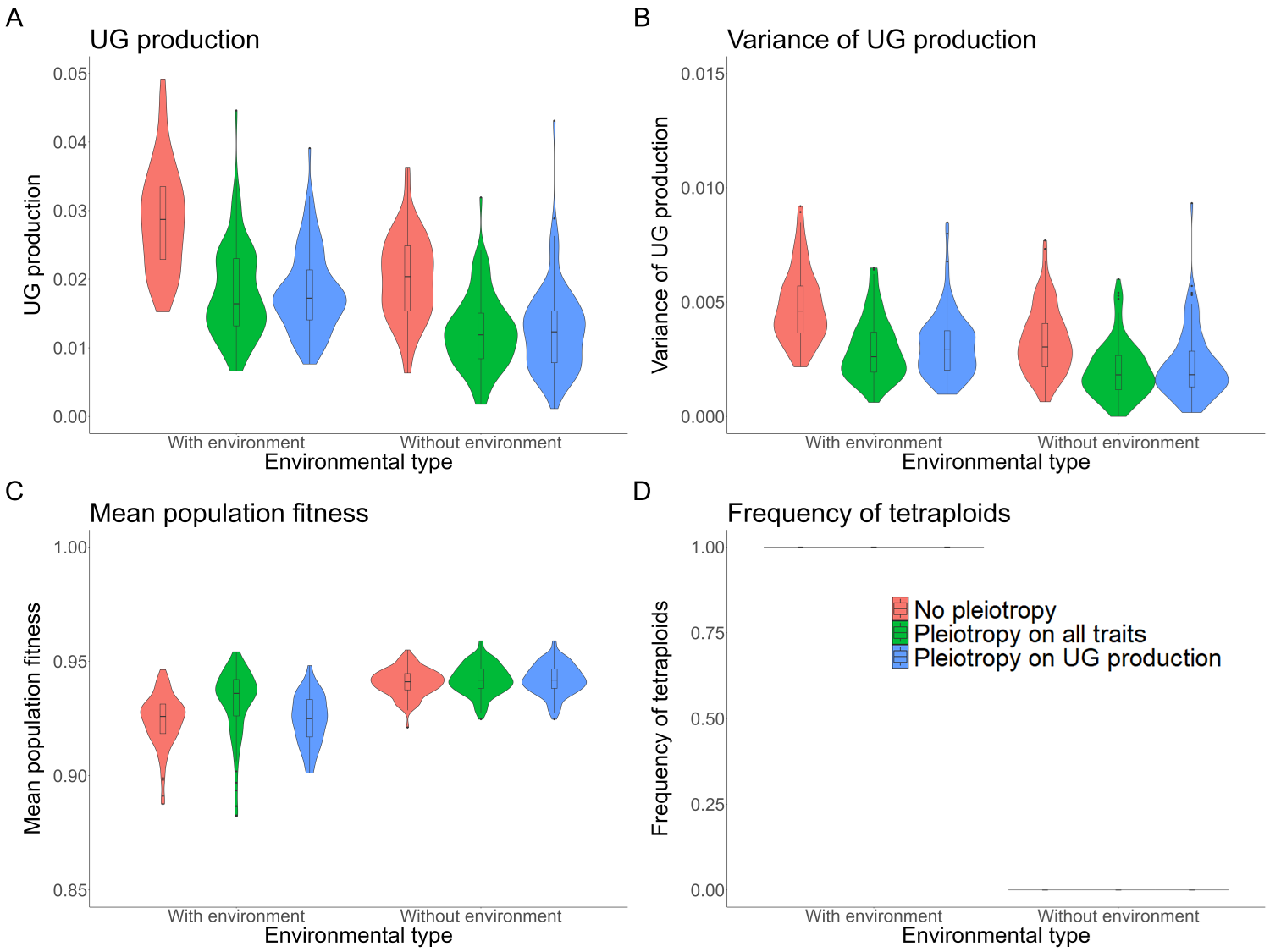

*Figure S3: The effect of Pleiotropy (number of pleiotropic loci = 80) type and environmental factor (0.3) on the unreduced gametes production and population fixation. (A) The unreduced gametes production of tetraploids with the 0.005 mutation rate; (B) The variation of unreduced gametes production with the 0.005 mutation rate. (C) The unreduced gametes production of tetraploids with the 0.005 mutation rate; (D) The variation of unreduced gametes production with the 0.005 mutation rate.*

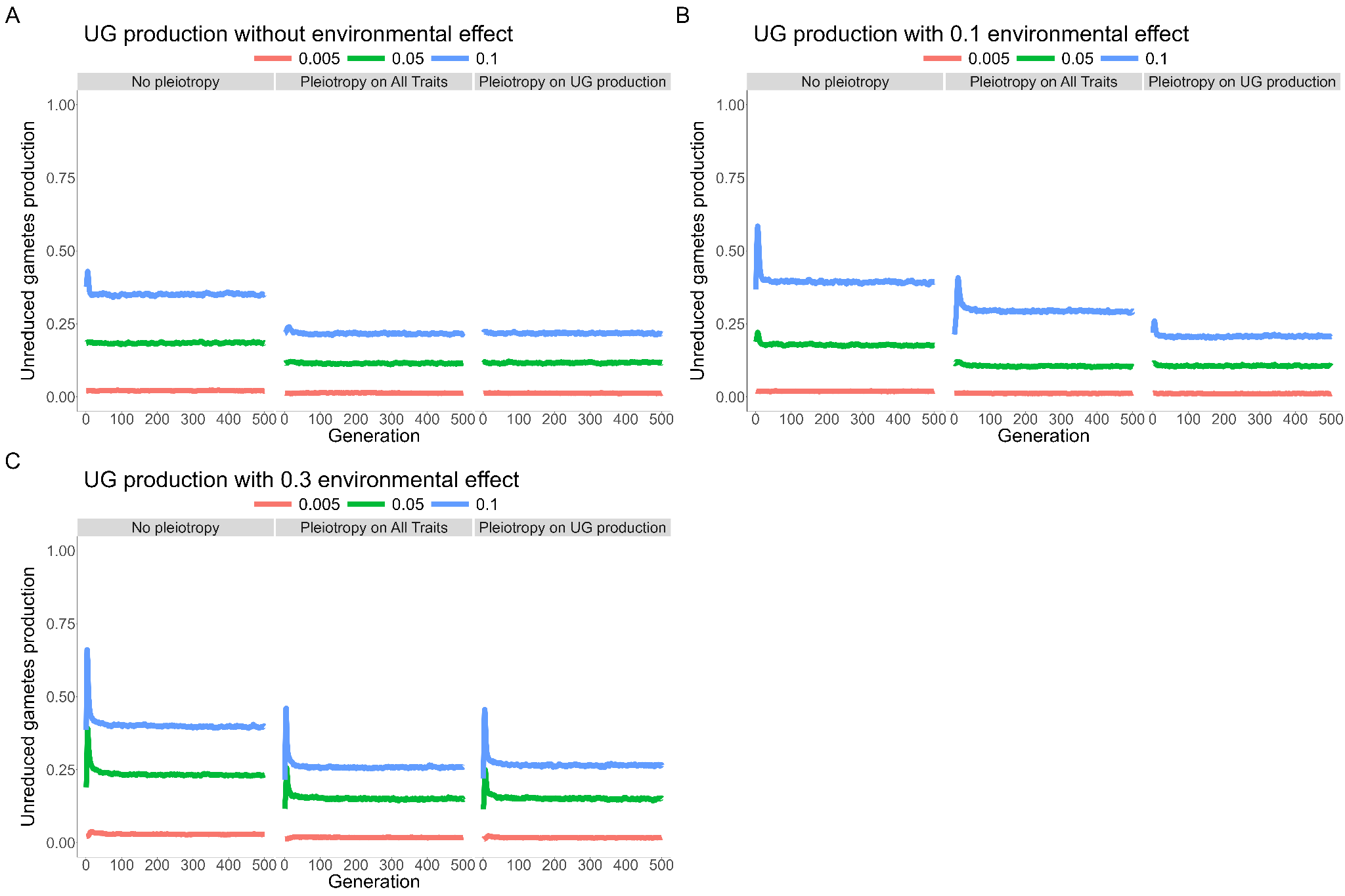

*Figure S4: Effect of pleiotropy type (no, all traits, unreduced gametes; number of pleiotropic loci = 80) and the environmental factor (0, 0.1, 0.3) on unreduced gametes production under different mutation rates (U = 0.005, 0.05, 0.1; differently colored lines), simulated from a population of genetically identical diploid individuals for 500 generations. Pleiotropy on all traits means that pleiotropy affected both the quantitative trait under directional selection and unreduced gametes production. (A) No environmental factor; (B) environmental factor 0.1; (C) environmental factor 0.3.*

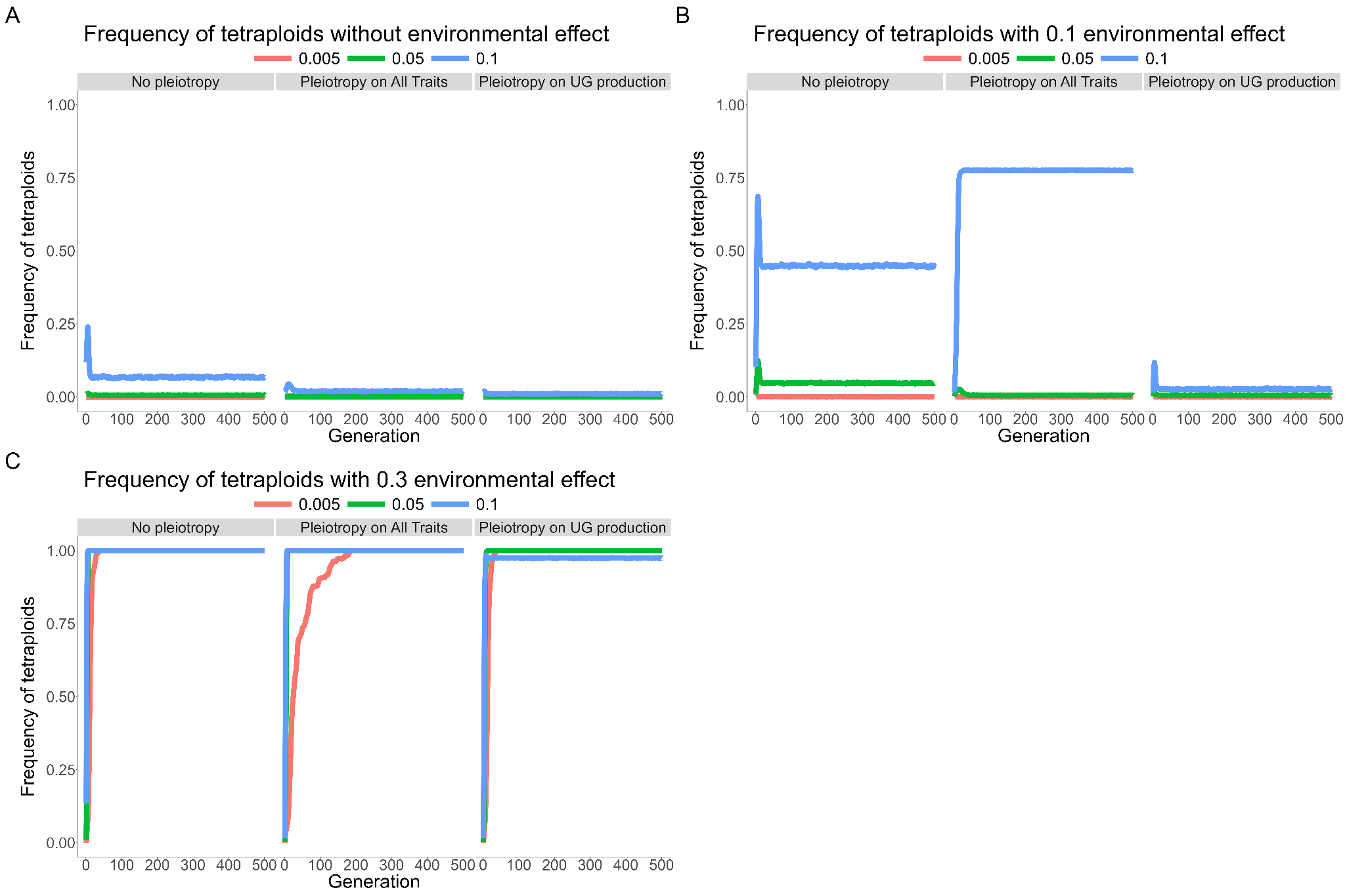

*Figure S5: Effect of pleiotropy type (no, all traits, unreduced gametes; number of pleiotropic loci = 80) and the environmental factor (0, 0.1, 0.3) on the frequency of tetraploids in the population under different mutation rates (U = 0.005, 0.05, 0.1; differently colored lines). simulated from a population of genetically identical diploid individuals for 500 generations. Pleiotropy on all traits means that pleiotropy affected both the quantitative trait under directional selection and unreduced gametes production. (A) No environmental factor; (B) environmental factor 0.1; (C) environmental factor 0.3.*
